# Density-dependent selection in *Drosophila*: evolution of egg size and hatching time

**DOI:** 10.1101/2021.10.24.465621

**Authors:** Srikant Venkitachalam, Srijan Das, Auroni Deep, Amitabh Joshi

## Abstract

Many different laboratory studies of adaptation to larval crowding in *Drosophila* spp. have all yielded the evolution of pre-adult competitive ability, even though the ecological context in which crowding was experienced varied across studies. However, the evolution of competitive ability was achieved through different suites of traits in studies wherein crowding was imposed in slightly different ways. Earlier studies showed the evolution of increased competitive ability via increased larval feeding rate and tolerance to nitrogenous waste, at the cost of food to biomass conversion efficiency. However, more recent studies, with crowding imposed at relatively low food levels, showed the evolution of competitive ability via decreased larval development time and body size, and an increase in the time efficiency of conversion of food to biomass, with no change in larval feeding rate or waste tolerance. Taken together, these studies have led to a more nuanced understanding of how the specific details of larval numbers, food amounts etc. can affect which traits evolve to confer increased competitive ability. Here, we report results from a study in which egg size and hatching time were assayed on three sets of populations adapted to larval crowding experienced in slightly different ways, as well as their low density ancestral control populations. Egg size and hatching time are traits that may provide larvae with initial advantages under crowding through increased starting larval size and a temporal head-start, respectively. In each set of populations adapted to some form of larval crowding, the evolution of longer and wider eggs was seen, compared to controls, thus making egg size the first consistent correlate of the evolution of increased larval competitive ability across *Drosophila* populations experiencing crowding in slightly different ways. Among the crowding-adapted populations, those crowded at the lowest overall eggs/food density, but the highest density of larvae in the feeding band, showed the largest eggs, on an average. All three sets of crowding-adapted populations showed shorter average egg hatching time than controls, but the difference was significant only in the case of populations experiencing the highest feeding band density. Our results underscore the importance of considering factors other than just eggs/food density when studying the evolution of competitive ability, as also the advantages of having multiple selection regimes within one experimental set up, allowing for a more nuanced understanding of the subtlety with which adaptive evolutionary trajectories can vary across even fairly similar selection regimes.

## Introduction

Populations adapted to high density conditions are expected to evolve greater competitive ability, a prediction highlighted by the theory of density-dependent selection, first formulated by MacArthur (1962) and MacArthur and Wilson (1967) (see Mueller 1997, 2009 for reviews on subsequent developments in this area). Several rigorous long-term selection experiments on populations of *Drosophila* reared under high larval density conditions subsequently validated this prediction, showing the evolution of increased pre-adult competitive ability in the crowding adapted populations when compared to their low density controls (Mueller 1988; Nagarajan *et al.* 2016; Sarangi *et al.* 2016). However, the traits that evolved as correlates of the increased pre-adult competitive ability differed widely across the studies (Nagarajan *et al.* 2016; Sarangi *et al.* 2016; Sarangi 2018).

The first of these experiments was done using two sets of replicate populations: the *K*-populations, maintained at high population density (larval and adult) by serial transfer, and the *r*-populations, maintained at low population density by culling (Mueller and Ayala 1981). Compared to the *r*-populations, the *K-* populations evolved greater larval competitive ability (Mueller 1988), increased larval feeding rate (Joshi and Mueller 1988), greater pupation height (Mueller and Sweet 1986; Joshi and Mueller 1993), greater larval foraging path length (Sokolowski *et al.* 1997), increased adult dry weight and pre-adult viability at high density (Bierbaum *et al.* 1989), and increased minimum larval food requirement for completion of development (Mueller 1990).

The next selection study sought to validate the results from the *r*- and *K*-populations, as the earlier selection regime confounded the effects of larval and adult crowding. Moreover, the *r*-populations were maintained on discrete generations, whereas the *K*-populations were maintained on overlapping generations (Mueller *et al.* 1993). Consequently, populations of *D. melanogaster*, originally derived from a different geographical region than the ancestors of the *r*- and *K*-populations, were used in a selection experiment that differentiated the effects of larval and adult crowding, and in which all selected populations and controls were maintained on a three-week discrete generation cycle (Mueller *et al.* 1993). The populations reared at high larval, but not adult, density were called the CU (Crowded as larvae, Uncrowded as adults), and the low density controls were called UU (Uncrowded as larvae, Uncrowded as adults) (Mueller *et al.* 1993). Similar to what was seen earlier in the *K*-populations, the CU populations evolved increased larval feeding rate and minimum larval food requirement for completion of development (Joshi and Mueller 1996), and larval foraging path length (Sokolowski *et al.* 1997). Moreover, the CU populations evolved increased pre-adult urea tolerance (Shiotsugu *et al.* 1997; Borash *et al.* 1998) and ammonia tolerance (Borash *et al.* 1998). The CU populations, however, did not evolve increased pupation height than controls (Joshi and Mueller 1996), unlike the *K*-populations; possible explanations are discussed by Joshi *et al.* (2003).

The broadly consistent results from the *r*- and *K*-populations and the CU and UU populations, together with similar results from the *rK* and *r×rK* populations (Guo *et al.* 1991), resulted in the canonical model for adaptation to larval crowding in *D*. *melanogaster* populations: these populations would exhibit increased pre-adult competitive ability and larval feeding rate, foraging path length, and tolerance to ammonia and urea, but would show reduced food to biomass conversion efficiency as a trade-off (Mueller 1997; Joshi *et al.* 2001; Prasad and Joshi 2003; Mueller *et al.* 2005; Mueller 2009; Mueller and Cabral 2012; Mueller and Barter 2015; Bitner *et al.* 2021). The canonical model was further strengthened by observations in *D*. *melanogaster* of greater pre-adult competitive ability in populations selected for increased larval feeding rate (Burnet *et al*. 1977), and the evolution of reduced pre-adult competitive ability in populations that evolved reduced larval feeding rate due to selection for either rapid pre-adult development (Prasad *et al*. 2001; Shakarad *et al.* 2005; Rajamani *et al.* 2006) or for increased parasitoid resistance (Fellowes *et al.* 1998, 1999).

The canonical model was, nevertheless, challenged later by three selection studies involving adaptation to larval crowding, in *D*. *ananassae, D. nasuta nasuta* and *D*. *melanogaster,* respectively (Nagarajan *et al.* 2016; Sarangi *et al.* 2016). In all three studies, crowding adapted populations did evolve greater larval competitive ability compared to their respective low density controls, but did so through a suite of traits different from the canonical model. No evolution of increased feeding rate was seen, nor were there any changes in urea tolerance, compared to controls. Instead, the crowding-adapted populations seemed to evolve greater larval competitive ability primarily through a decrease in pre-adult development time, expressed even when assayed at low density, and an increase in the time efficiency of food to biomass conversion, relative to controls (Nagarajan *et al.* 2016; Sarangi *et al.* 2016). It then became apparent that the major difference between these studies and the earlier work that had given rise to the canonical model was in the ecological details of the context in which larvae in selected populations experienced crowding (Sarangi 2018). Specifically, the populations used by Nagarajan *et al.* (2016) and Sarangi *et al.* (2016) had very low amounts of food per vial, whereas the earlier studies had used larger amounts of food and a greater number of eggs. For example, the MCU populations of Sarangi *et al.* (2016) were maintained at a density of about 600 eggs per vial containing 1.5 mL food whereas the CU populations (Mueller *et al.* 1993) were reared in vials containing about 1500 eggs in 6-7 mL of food. Subsequently, altering the amount of food and number of eggs while keeping overall eggs per unit food density the same was shown to affect pre-adult survivorship and development time, as well as the weight distribution of eclosing flies (Sarangi 2018). Therefore, in order to examine this phenomenon further, two new sets of *D*. *melanogaster* populations were subjected to selection for adaptation to larval crowding. One set of four populations, called LCU, was maintained at around 1200 eggs in 6 mL food, and this regime was meant to approximate the CU populations of Mueller *et al.* (1993). The other set of four populations was called the CCU, and was maintained at twice the number of eggs and twice the volume of food, and thus an identical overall density, as the MCU populations (Sarangi 2018). Thus, a system of 16 populations was created: ancestral controls (MB), MCU, CCU and LCU, with four replicate populations in each regime (Sarangi 2018).

Interestingly, although the LCU and CCU populations did evolve greater pre-adult competitive ability compared to the MB controls (Sarangi 2018; S. Venkitachalam and A. Joshi, *unpubl. data*), they did so via an increased larval feeding rate, unlike the MCU populations (Sarangi 2018). However, as in the MCU populations, no evolution of pre-adult urea or ammonia tolerance was seen in the CCU and LCU populations (Sarangi 2018). The overall picture that emerges is, thus, one of ‘unity in ends, diversity in means’, with even populations experiencing identical larval density in slightly different ecological contexts exhibiting the evolution of increased pre-adult competitive ability with or without a concomitant increase in larval feeding rate (Sarangi 2018). Here, we show that there is nevertheless a commonality in evolutionary trajectories across the MCU, CCU and LCU populations in that they all seem to have evolved a shorter egg hatching time and a greater egg size than the MB controls. These traits may be important for larval competitive ability, as together they can effectively provide a temporal head-start and initial size advantage in competition (Sokolowski *et al.* 1997; Bakker 1961, 1969).

## Materials and methods

### Experimental populations

We used four sets of long-term laboratory populations of *D. melanogaster*, with each set consisting of four replicate populations, as briefly described below. The derivation and maintenance of all these populations have been discussed in detail by Sarangi (2018).

#### MB 1-4

These are four low density reared populations that serve as ancestral controls to the three sets of crowding-adapted populations. They are maintained at a relatively low density of approx. 70 eggs in 6 mL of cornmeal-sugar-yeast medium, in cylindrical Borosilicate glass vials of 2.2-2.4 cm inner diameter and 9.5 cm height.

#### MCU1-4

These populations experience larval crowding at ~600 eggs in ~1.5 mL of cornmeal medium, in the same type of vials as MBs. At the time of assaying, the MCUs had undergone at least 218 generations of selection (blocks (i.e. replicate populations) 1, 2 assayed at gen. 218; blocks 3, 4 assayed at gen. 219).

#### CCU1-4

These populations experience larval crowding at ~1200 eggs in ~3 mL of cornmeal medium, in the same type of vials as MBs. It should be noted that MCU and CCU have the exact same overall eggs/food density. At the time of assaying, the CCUs had undergone at least 97 generations of selection (blocks 1, 2 assayed at gen. 97; blocks 3, 4 assayed at gen. 98).

#### LCU 1-4

These populations experience larval crowding at ~1200 eggs in ~6 mL of cornmeal medium, in Borosilicate glass vials of ~2 cm inner diameter and ~9 cm height (approx. 6-dram volume, to mimic the CU populations of Mueller *et al.* (1993)). At the time of assaying, the LCUs had undergone at least 96 generations of selection (blocks 1, 2 assayed at gen. 96; blocks 3, 4 assayed at gen. 97).

While the pre-adult stages of each population are maintained in vials, the adults are transferred to Plexiglas cages (25 × 20 × 15 cm^3^) on the day of eclosion. Given the low larval density of MB populations, they are transferred to cages on the 11^th^ day from egg collection. In the crowding-adapted populations, there is a large amount of variation in eclosion time and thus, transfer of eclosing adults to cages is done daily from day 8 to day 21 from egg collection. Fresh cornmeal food plates are given (following a fresh plate given on initiation of transfers) on day 10, 12, 14 and 17 from egg collection. On day 18 from egg collection, the flies in the cages are provided a food plate with a generous smear of a paste of live yeast mixed with water and a few drops of glacial acetic acid. On day 20 from egg collection, the flies are provided cornmeal food plates with vertical edges present for egg laying, for 18 hours. Finally, eggs laid by the flies on these plates are used to initiate the next generation, with eggs being transferred to fresh vials containing the respective food volume assigned to each population. All populations are maintained under constant light, at 25 ± 1°C and 70-90% relative humidity.

### Standardisation of populations

Prior to assays, all populations were subjected to one generation of standardisation (rearing in a common low larval density environment), to eliminate any non-genetic parental effects. Eggs from each population were collected at approx. 70 eggs in 6 mL of food per vial, for a total of 40 vials per population. The flies eclosing in these vials were transferred to cages on day 11 from egg collection, following which they were provided a food plate, with a generous smear of the live yeast-water-acetic acid paste, for approx. 48 hours. On day 13 from egg collection, the flies were provided a food plate for egg collection for around 18 hours, and two rearing environments for the assay were set up on day 14 from egg collection. All assays were conducted in constant light, at 25 ± 1°C and 70-90% relative humidity.

### Rearing environments

For each population, the eggs collected from the previous standardised generation were used to form two sets of assay populations, reared at two larval densities.

#### Low density rearing

The first set was kept at a relatively low eggs/food density of ~70 eggs in 6 mL cornmeal medium per vial, with a total of 40 vials per replicate population. As in the standardisation, the adults eclosing in the vials were transferred to a cage on day 11 from egg collection. On day 17 from egg collection, the flies were provided a food plate with a generous smear of live yeast paste for ~48 hours. Following this, a “dummy” egg collection cornmeal plate was provided for an hour, which was for the laying of any eggs previously incubating inside the females. Relatively synchronized ggs for the hatching time and egg size assays were then obtained by providing a harder plate with double the usual agar and different composition (only yeast, sugar added), for 45 minutes. This composition ensured easier egg removal for counting.

#### High-density rearing

Eggs for the second set were collected into vials at a relatively high eggs/food density – approx. 300 eggs in 2 mL cornmeal medium per vial, with a total of 12 vials per population. This simple density change was done as a first pass to obtain reduced adult size without impacting survivorship greatly. Unlike in the low density rearing conditions, adults emerging from the vials were transferred to cages daily from the day of the start until the end of eclosion, usually day 15-16. The protocols from day 17 onwards were the same as in the low density reared populations.

### Egg hatching time

The assay was carried out in plastic Petri plates (90 mm diameter × 14 mm height), in which a thin layer of 12 g/L agar solution (containing 2.4 g/L methyl 4-hydroxybenzoate, as preservative) was spread. A 6 × 6 square grid (36 square cells, each having 3 mm sides) was pasted on the bottom of each Petri plate, which was visible through the transparent layer of agar. A total of 5 Petri plates were used per selection × rearing density × block combination, with 36 eggs per Petri plate – one egg per cell of the grid (Figure 1). Checks for egg hatching were done at 13, 15, 17, 18, 19, 20, 21, 22, 24, 26, 28 and 30 hours from egg laying, respectively. At each check, eggs which hatched in the time interval between the current and previous check were noted.

**Figure 1:**
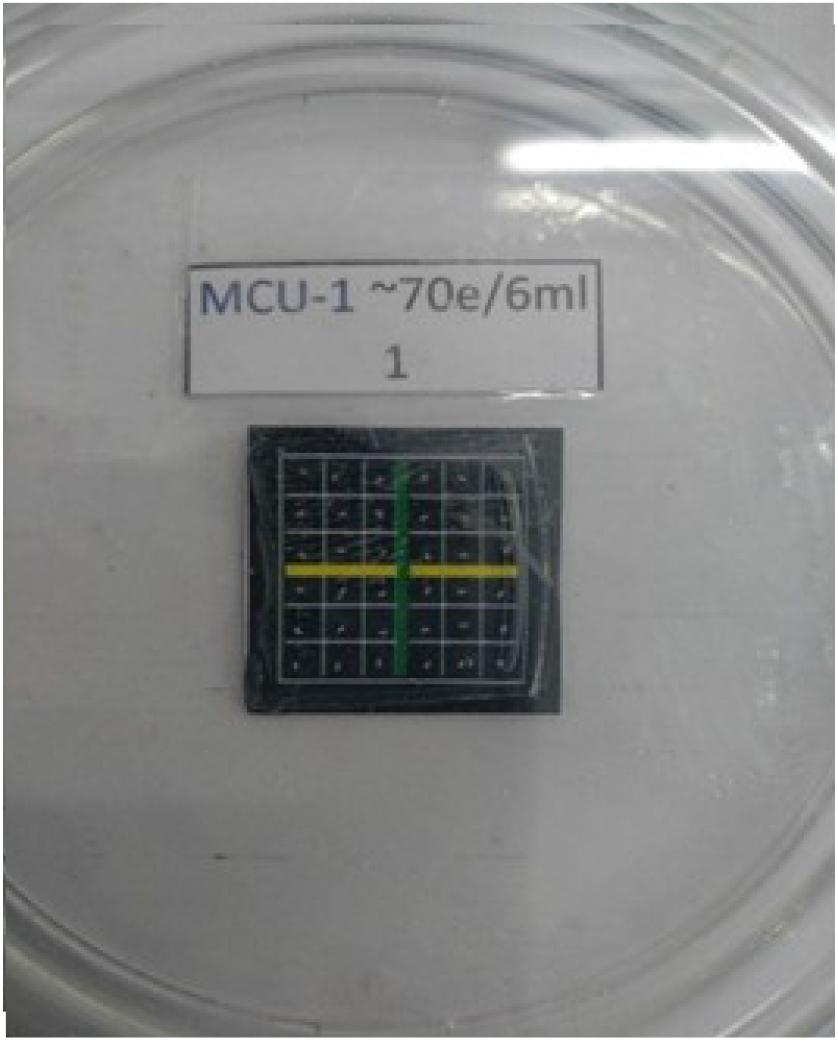
Apparatus for egg hatching time and hatchability measurements. The 6 × 6 grid is pasted on the bottom of a Petri plate containing a thin layer of agar solution. Each cell of the grid contains an egg, as can be seen in the image. The label denotes the selection × rearing density × block combination used, along with the replicate plate number (‘1’ in this case).

### Egg hatchability

From the hatching time assay, we also recorded how many eggs hatched within 48 hours from egg laying were noted. The egg hatchability was calculated as the number of eggs hatched divided by the total number of eggs. Earlier hatchability experiments on populations with relatively close ancestry to our MB populations did not use clear eggs due to their infertility (Chippindale *et al.* 1997). However, we have found that some clear eggs in our populations can lead to viable adults (S. Venkitachalam, *pers. obs*.), and thus we used all but the visibly damaged eggs for our experiments.

### Egg length and width

For size measurements, a total of 30 eggs (obtained as 10 eggs each in 3 replicates) were measured per selection × rearing density × block combination. The eggs were placed on a Neubauer haemocytometer and photographed under a stereo-microscope. The parallel lines on the haemocytometer, which were a known distance apart (200 μm or 250 μm, depending on the set of lines used; see Figure 2) provided a scale with which to measure the eggs. Egg length (estimate of polar axis) and egg width (estimate of minor axis) were measured from the photographs (Figure 2) using ImageJ (Rasband 1997-2018).

**Figure 2:**
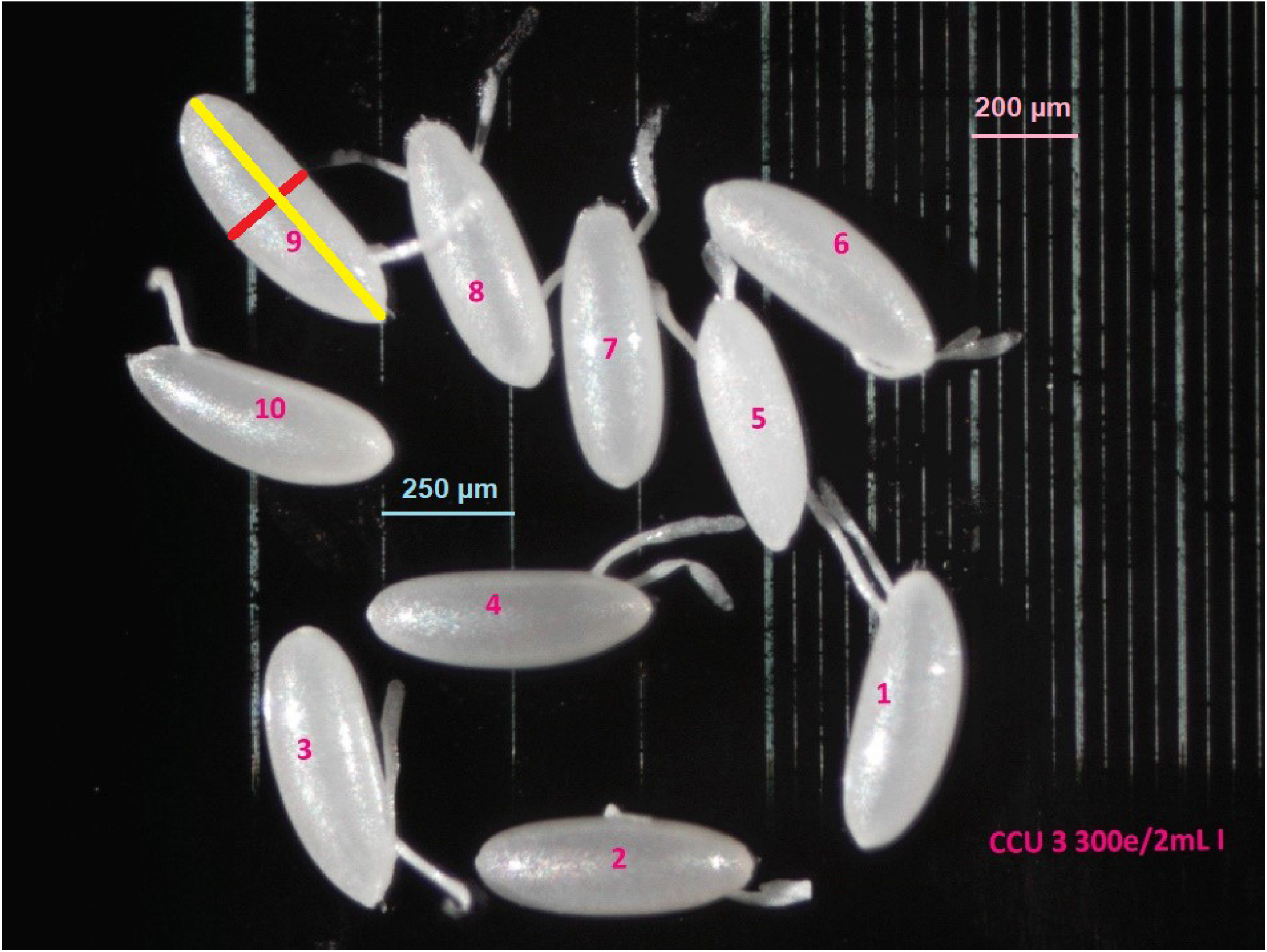
Egg size measurement setup for a replicate containing 10 eggs. There were three such replicates for each selection × rearing density × block combination. Eggs were numbered from 1 through 10. The egg labelled ‘9’ has two lines of measurement drawn for demonstration: the yellow line denotes egg length and the red line, egg width. The background is that of a Neubauer haemocytometer, which contains parallel lines set a known distance apart, and can thus be used to determine the scale in the image (parallel lines set either 250 μm or 200 μm apart could be used, as marked in the figure).

### Statistical analyses

Every replicate larval crowding adapted population shares ancestry with an MB population with the same replicate subscript i.e. replicate population *i* in the MCU, CCU and LCU regimes is derived from replicate *i* of MB (*i* = 1..4). This permits the use of a completely randomized block design in our statistical analysis, with replicate populations bearing the same subscript treated as blocks. Assays were conducted concurrently on all populations of a block. The data were subjected to a mixed model ANOVA (type III) in a fully factorial design, with the block (4 levels) treated as a random factor. Selection (4 levels) and rearing density (2 levels) were treated as fixed factors. For hatchability, the analysis was repeated after performing an arcsine square root transformation on the data, to check for differences in the statistical significance of the fixed factors. All ANOVAs were done using STATISTICA™ Windows release 5.0 (Statsoft 1995). Tukey’s HSD was used for post-hoc pairwise comparisons at α = 0.05. The image measurements for egg size were done using ImageJ (Rasband 1997-2018). Pearson’s product-moment correlation coefficients were calculated pairwise for population means of egg hatching time, egg length and egg width.

## Results

### Egg hatching time

Mean egg hatching time across all four types of selected and control populations was close to 20 hours, and the range of variation among means was only about 30 min (Figure 3). However, all three sets of selected populations had shorter mean hatching times than the MB controls, the shortest being in the LCU populations, followed by CCU, and then by MCU (Figure 3). The ANOVA revealed a significant main effect of selection (*F*_3,9_ = 4.142, *P* = 0.042) on egg hatching time, but post-hoc pairwise comparisons showed a significant difference only between LCU and MB (Figure 3). There neither a significant main effect of rearing density (*F*_1,3_ = 0.631, *P* = 0.485), nor a significant selection × rearing density interaction (*F*_3,9_ = 1.871, *P* = 0.205).

**Figure 3:**
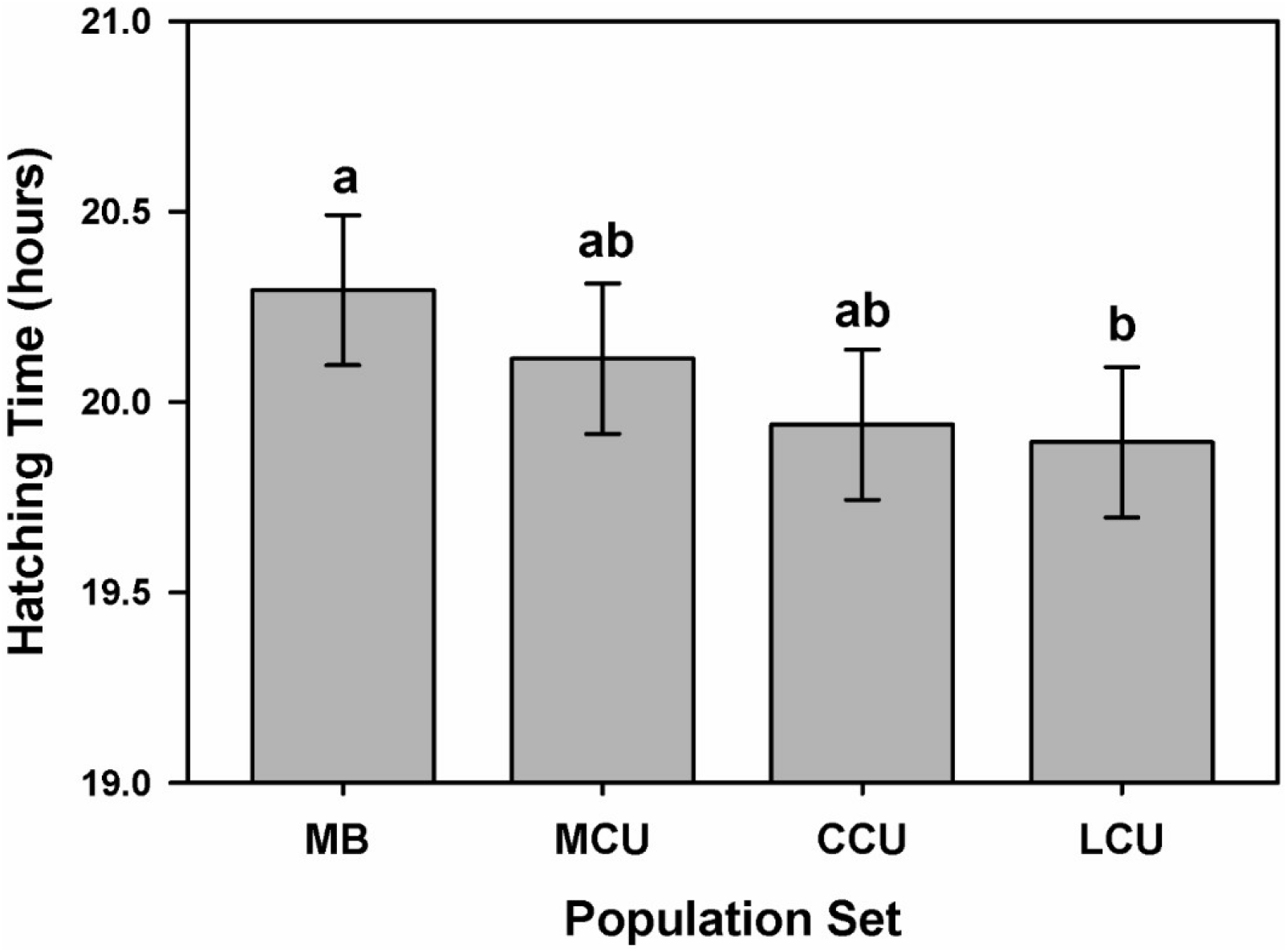
Mean egg hatching time in hours for the four levels of selection, averaged over all levels of rearing density and block. The error bars show 95% confidence intervals, calculated from post-hoc Tukey’s HSD, and allow for visual hypothesis testing – identical superscript letters denote means that did not differ significantly, whereas different letters denote means that differed significantly.

### Egg hatchability

Mean egg hatchability ranged from about 75-90% across selection × rearing density combinations, with flies reared as larvae at high density (300 eggs in 2 mL food) tending to lay more viable eggs than those reared at low density (70 eggs in 6 mL food), most markedly so in the MCU populations (Figure 4). The ANOVA revealed no significant main effect of selection (*F*_3,9_ = 1.067, *P* = 0.410). There was, however, a significant main effect of rearing density (*F*_1,3_ = 12.484, *P* = 0.039), as well as a significant selection × rearing density interaction (*F*_3,9_ = 4.236, *P* = 0.040). Post-hoc comparisons revealed that only MCU showed significantly higher hatchability when flies were reared as larvae at high versus low density. Similar but non-significant differences were also seen in the MB and LCU, whereas mean hatchability of CCU reared at low versus high larval density was very similar (Figure 4). The pattern of significant ANOVA effects was unaffected by whether untransformed or arcsine transformed data were used.

**Figure 4:**
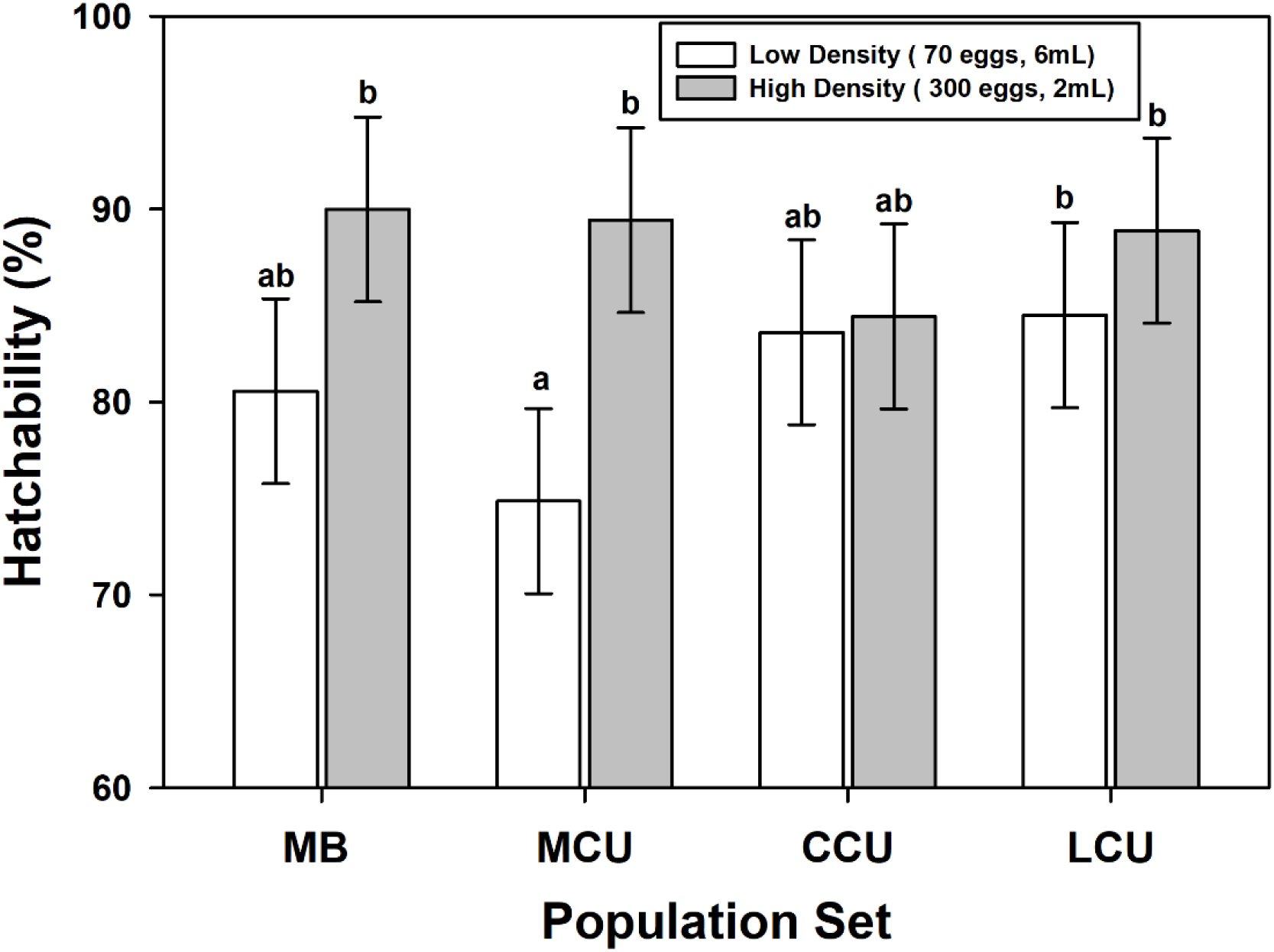
Mean hatchability (%), for all combinations of four levels of selection and two levels of rearing density, averaged across all blocks. The error bars show 95% confidence intervals, calculated from post-hoc Tukey’s HSD, and allow for visual hypothesis testing – identical superscript letters denote means that did not differ significantly, whereas different letters denote means that differed significantly.

### Egg length (μm)

Mean egg length in MB populations was significantly less than any of the sets of populations selected for larval crowding (Figure 5), driving a significant ANOVA main effect of selection (*F*_3,9_ = 22.104, *P* < 0.001). Egg length, on an average, did not differ significantly between rearing densities (main effect of rearing density: *F*_1,3_ = 8.109, *P* = 0.065; Figure 5). LCU eggs were longer than those of MCU across both rearing densities, but longer than CCU eggs only at high rearing density. On the other hand, MCU eggs were shorter than CCU eggs at low density, but of similar length at high density (Figure 5), and these rearing densityspecific differences among various crowding adapted sets of populations drove a significant ANOVA selection × rearing density interaction (*F*_3,9_ = 4.830, *P* = 0.029). This pattern of differences between CCU and the other crowding adapted populations was likely due to an average of 9 μm longer eggs laid by CCU females when reared at low as compared to high density, although this difference itself was not statistically significant (Figure 5).

**Figure 5:**
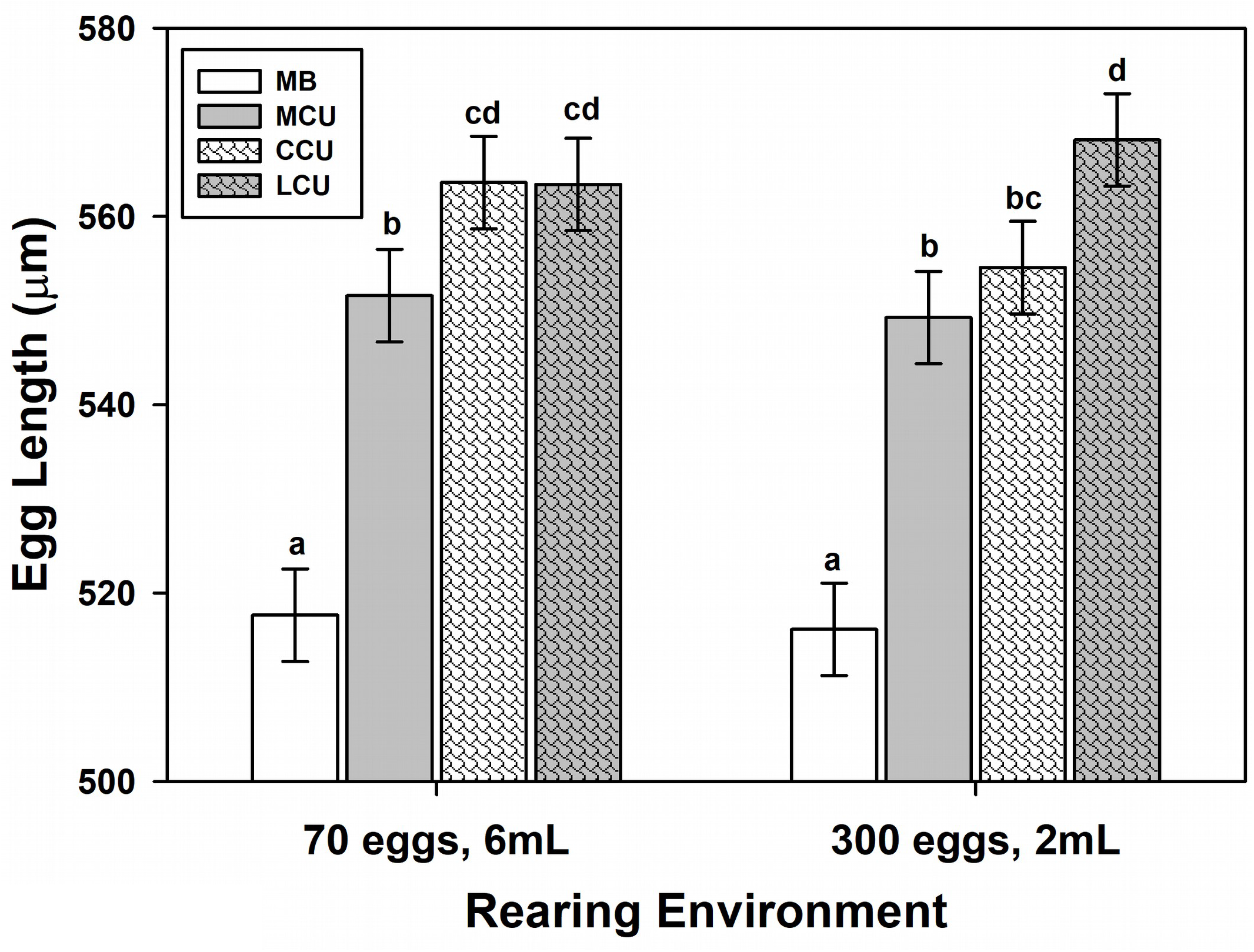
Mean egg length (μm), for all combinations of four levels of selection and two levels of rearing density, averaged across all blocks. The error bars show 95% confidence intervals, calculated from post-hoc Tukey’s HSD, and allow for visual hypothesis testing – identical superscript letters denote means that did not differ significantly, whereas different letters denote means that differed significantly.

### Egg width (μm)

Overall, the egg width data were fairly similar to those for egg length (Figures 5,6), with egg width being considerably lower in MB populations compared to all crowding adapted populations (ANOVA main effect of selection: *F*_3,9_ = 5.496, *P* = 0.020), and not differing, on an average, between rearing densities (main effect of rearing density: *F*_1,3_ = 0.584, *P* = 0.500) (Figure 6). Eggs laid by MCU, CCU and LCU flies reared at low density did not differ much in mean width, whereas at high rearing density LCU females laid the widest eggs, and the MCU and CCU did not significantly differ in egg width (selection × rearing density interaction: *F*_3,9_ = 7.971, *P* = 0.007; Figure 6).

**Figure 6:**
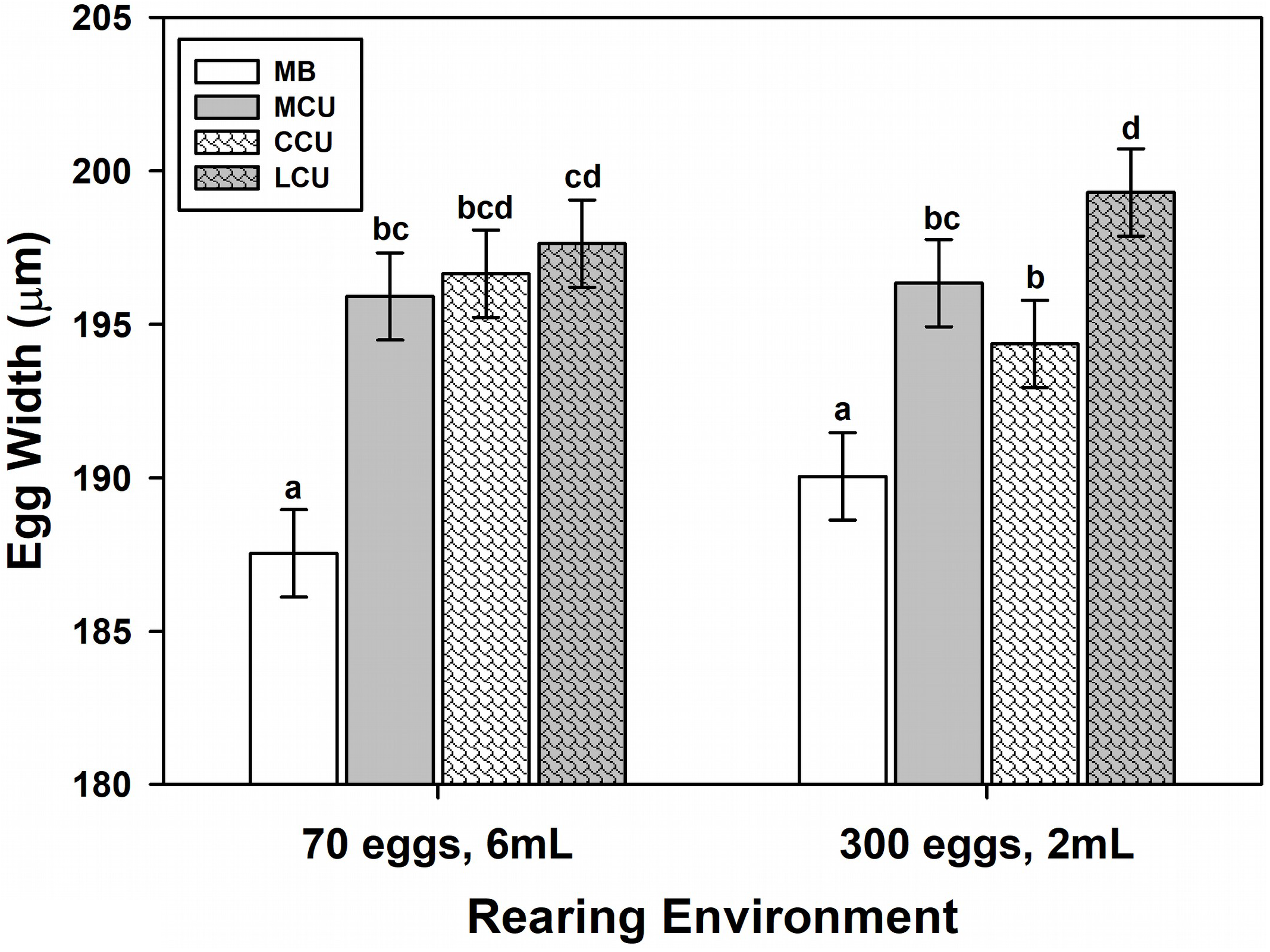
Mean egg width (μm), for all combinations of four levels of selection and two levels of rearing density, averaged across all blocks. The error bars show 95% confidence intervals, calculated from post-hoc Tukey’s HSD, and allow for visual hypothesis testing – identical superscript letters denote means that did not differ significantly, whereas different letters denote means that differed significantly.

### Trait correlations

‘There was a strong, positive correlation across population means between egg length and width (*r* = +0.771, *P* < 0.001; Figure 7), indicating that populations with longer eggs also tended to have wider eggs, and vice versa. The correlations for mean hatching time and mean egg length (*r* = −0.395, *P* = 0.025), and for mean hatching time and mean egg width (*r* = −0.548, *P* = 0.001) were both negative, although the strength of the correlation was moderate in both cases, being stronger for hatching time with egg width (Figure 7). There were no discernible patterns for within population correlations between egg length and egg width, with the mean correlation coefficient being around 0.11, and no selection × rearing density combination exceeding a correlation coefficient of 0.3 (data not shown).

**Figure 7:**
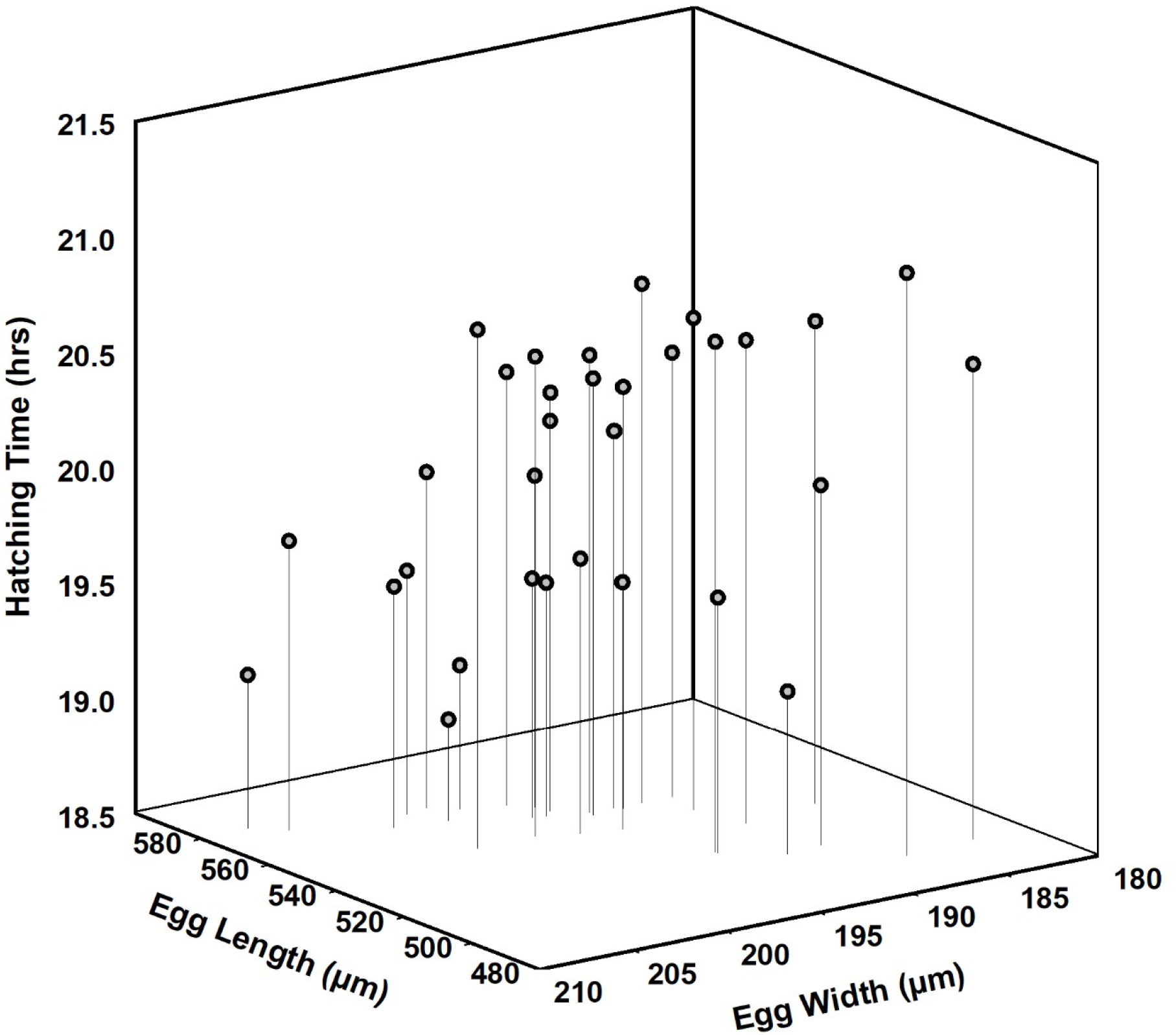
The relationship between mean egg length (μm), mean egg width (μm) and mean hatching time (hours) across the four sets of populations. Each data point represents the mean trait value for the three traits in one combination of selection × rearing density × block. Note the orientation of the x and y axes.

## Discussion

Despite the variation in which traits underlie the evolution of greater pre-adult competitive ability in *Drosophila* populations that experience larval crowding under slightly varying conditions (reviewed in Sarangi 2018), our results suggest one common adaptation across at least three such selection regimes covering a range of egg number and food amount combinations that more or less mimics the range of previous studies. Adults from the MCU, CCU and LCU populations laid eggs with greater length and width compared to the MB populations, when assayed at low (70 eggs in 6 mL food) or relatively high (300 eggs in 2 mL) density rearing conditions (Figures 5 and 6). Along with the strong positive correlation seen between the mean egg length and mean egg width across populations (Figure 7), these results indicate an increase in overall egg size of all these three sets of crowding adapted populations compared to the ancestral controls. Our results are also in agreement with earlier study from a different laboratory, which demonstrated an increase in egg size, relative to controls, in crowding adapted populations derived from our MCU populations and maintained on a similar regime (Kumar 2014).

The eggs laid by LCU females were larger than those laid by MCU at both low and high density rearing conditions, with CCU eggs being intermediate in size (Figures 5 and 6). The differences among the MCU, CCU and LCU populations themselves are perhaps just as important as the consistent difference between the egg size of the crowding-adapted and MB populations. While previous comparisons of results from selection studies in differently crowded cultures have focused on the repeatability of qualitative differences found between a single set of crowding adapted populations against its controls (Joshi and Mueller 1996; Nagarajan *et al.* 2016; Sarangi *et al.* 2016), our study system permits more nuanced, quantitative comparisons between multiple types of high-density selection regimes.

The importance of plasticity in egg size has been studied extensively from the perspective of non-genetic maternal effects, in the contexts of both competition and malnutrition (Kawecki 1995; Azevedo *et al.* 1997; Prasad *et al.* 2003; Vijendravarma *et al.* 2010; Yanagi *et al.* 2013). In our study, egg size did not show any statistically significant difference between parents reared at low or high larval density (Figures 5,6). However, the CCU populations did show a consistent trend for smaller eggs when crowded at the given density. It may be possible that either the relative scaling of egg size with female body size, or the sensitivity of female size to larval crowding, might be different across the MCU, CCU and LCU populations. This might be worth exploring in the future.

Moreover, since it is also known that crowding more severe than what we used can further decrease body size (Sang 1949; Bakker 1961; S. Venkitachalam and A. Joshi *unpubl. data*), observed effects of rearing density on egg traits in these populations may change under more extreme crowding, where size at eclosion and pre-adult survivorship are more severely impacted than they were in this study.

If we compare our egg size results with those obtained in a comparison of populations selected for rapid preadult development (FEJ) relative to their controls (JB), which are similar to the MB populations (first described in Prasad *et al*. 2000), there are some interesting similarities and differences. Although the MCU, CCU and LCU populations all have reduced pre-adult development time compared to MB controls (Sarangi 2018), the FEJ populations had undergone a far greater reduction in pre-adult development time, relative to their controls, as that was the primary trait under selection (Prasad et al. 2000; Prasad and Joshi 2003). On an average, eggs laid by FEJ females, after rearing at low density as larvae, were 3.8% longer, 7% wider, and 11% heavier than those of their controls (B.M. Prakash and A. Joshi, *unpubl. data*). The MCU, CCU and LCU populations in this study exhibited length increases of 6.5%, 8.2% and 9.5%, respectively, compared to the MB controls, and the corresponding width increases were 3.9%, 3.6% and 5.1%. From this comparison, we might conclude that MCU, CCU and LCU eggs are likely to be about 10-15% heavier than MB eggs. Interestingly, in the FEJ populations, the increase in width was greater than in length; it is just the opposite in the MCU, CCU and LCU populations. At this point, we cannot say why this may be so, although, given the very different selection pressures (rapid development vs. larval crowding), the mechanisms underlying the response could differ. The difference is unlikely to be explained by female size differences, since flies of FEJ, as well as MCU, CCU and LCU populations tend to be quite small relative to controls.

Although eggs from all crowding-adapted populations hatched faster than those of the controls, only the difference between mean egg hatching time between the LCU and MB populations was statistically significant (Figure 3). Moreover, the difference between LCU and MB mean egg hatching time was only ~30 minutes. However, given the egg size results (Figures 5,6), the pattern of MB > MCU > CCU > LCU for egg hatching time (Figure 3), and the negative correlation between mean egg length and mean hatching time, as well as between mean egg width and mean hatching time, we might expect crowding adapted populations that evolve increased egg size and decreased hatching time to benefit from a potent head-start in conditions of high pre-adult competition. Thus, we might expect LCU larvae to have a greater head-start in terms of pre-adult competition, compared to MCU larvae, and much greater still compared to MB larvae. This does not, however, necessarily imply that LCU larvae will have greater pre-adult competitive ability than MCU larvae, as differences in growth rates, efficiency and waste tolerance may also play a major role in determining pre-adult competitive ability (Bakker 1961; Joshi and Mueller 1996; Santos *et al.* 1997; Borash *et al.* 1998; Nagarajan *et al.* 2016; Sarangi *et al.* 2016). The evolution of a greater potential headstart in the LCU populations could be driven by the fact that, compared to the MCU and CCU populations, the LCU larvae experience the highest density within the feeding band (the few mm deep zone below the food surface within which larvae feed), even though their overall eggs/food density is lower than that in the other two selection regimes.

Overall hatchability was lower in our study than usually observed in related populations (e.g. over 90% in Chippindale *et al.* (1994)). This might be attributed to reduced humidity due to the very thin layer of agar used by us – future experiments using a thicker agar layer or regular cornmeal food might alleviate the survivorship, if this explanation is correct. We also observed reduced hatchability of eggs laid by MCU flies reared under low density conditions (Figure 4). It is not clear if this is due to an increase in infertile or unviable eggs (Chippindale *et al.* 1994, 1997), and whether it is driven by some correlated response(s) to evolution under larval crowding for over 200 generations of selection in the MCU populations, much longer than their CCU and LCU counterparts.

In conclusion, our results highlight increased egg size as being a consistent evolutionary correlate of greater pre-adult competitive ability across three differently crowded selection regimes that otherwise differ in the traits they have evolved in response to chronic larval crowding. Moreover, adults from populations crowded with the lowest eggs/food density, but the highest feeding band density, laid the largest eggs with the fastest hatching times, thus potentially allowing for a substantial head-start in the context of pre-adult competition. The study system we describe allows the comparison of adaptations to different crowding scenarios, highlighting quantitative differences that may otherwise not be possible to see when comparing qualitative results between different long-term selection experiments.

## Acknowledgments

We thank Sajith V. S., Ramesh Kokile, Avani Mital, Medha Rao, Bhavya Pratap Singh, Rajanna N. and Muniraju P. for help with the experiments. S. Venkitachalam was supported by a doctoral fellowship from the Jawaharlal Nehru Centre for Advanced Scientific Research. S. Das and A. Deep had their stay supported by the Jawaharlal Nehru Centre for Advanced Scientific Research’s Project Oriented Biological Education and Summer Research Fellowship Programme, respectively. This work was supported by a J. C. Bose National Fellowship from the Science and Engineering Research Board, Government of India, to A. Joshi and in part by A. Joshi’s personal funds.

## Notes

### Competing Interest Statement

The authors have declared no competing interest.

